# How community adaptation affects biodiversity-ecosystem functioning relationships

**DOI:** 10.1101/867820

**Authors:** Flora Aubree, Patrice David, Philippe Jarne, Michel Loreau, Nicolas Mouquet, Vincent Calcagno

## Abstract

Evidence is growing that evolutionary dynamics can impact biodiversity-ecosystem functioning (BEF) relationships. However the nature of such impacts remains poorly understood. Here we use a modelling approach to compare random communities, with no trait evolutionary fine-tuning, and co-adapted communities, where traits have co-evolved, in terms of emerging biodiversity-productivity, biodiversity-stability, and biodiversity-invasion relationships. Community adaptation impacted most BEF relationships, sometimes inverting the slope of the relationship compared to random communities. Biodiversity-productivity relationships were generally less positive among co-adapted communities, with reduced contribution of sampling effects. The effect of community-adaptation, though modest regarding invasion resistance, was striking regarding invasion tolerance: co-adapted communities could remain very tolerant to invasions even at high diversity. BEF relationships are thus contingent on the history of ecosystems and their degree of community adaptation. Short-term experiments and observations following recent changes may not be safely extrapolated into the future, once eco-evolutionary feedbacks have taken place.

## Introduction

Diversity, most classically defined as the number of constituent species in a community, plays an essential role in many aspects of ecosystem functioning (Hooper *et al.*, 2005, 2012; Isbell *et al.*, 2011). Understanding how species composition affects ecosystem properties is a fundamental question in basic and applied ecology, and renewed practical importance given the accelerating biodiversity crisis (Pimm *et al.*, 2014).

Observational data, controlled experiments and theoretical developments have converged in identifying ecosystem properties that exhibit systematic responses to diversity. Three types of so-called biodiversity-ecosystem functioning (BEF) relationships are most commonly described, even though all three are seldom considered in the same study. (i) First, the biodiversity-productivity relationship, historically investigated in grassland communities (Tilman *et al.*, 1996; Loreau & Hector, 2001), has been explored in several other taxa and ecosystems (Abramsky & Rosenzweig, 1984; Naeem *et al.*, 1994; Hooper *et al.*, 2005; Gamfeldt *et al.*, 2015). It is often assumed that more diverse ecosystems are more productive, in agreement with theoretical predictions (Loreau, 1998; Tilman, 1999). (ii) Second, biodiversity-stability relationships have also received a lot of attention (Elton, 1958; Tilman, 1999; McCann, 2000), both theoretically (May, 1973; Loreau & Mazancourt, 2013) and experimentally (Gross *et al.*, 2014; Renard & Tilman, 2019). The intuitive view that diverse ecosystems are more stable in the face of environmental fluctuations appeared contradicted by early theoretical models suggesting the opposite (McCann, 2000). In fact, predictions may differ importantly depending on the type of stability metric, with negative relationships expected at the level of individual species (dynamical stability: May, 1973; Tilman *et al.*, 1996; Ives & Carpenter, 2007), and positive relationships expected for aggregate metrics (ecosystem stability: May, 1973; Tilman *et al.*, 1996; Ives *et al.*, 1999; Barabás & D’Andrea, 2016; Pennekamp *et al.*, 2018). (iii) Last, biodiversity-invasion relationships have also attracted much attention, since native diversity has long been regarded as a key attribute determining the susceptibility of communities to invasions. It is generally considered that more diverse ecosystems should be less susceptible to invasions, and should suffer from fewer adverse impacts (e.g. secondary extinctions) following an invasion (Levine, 2000; Hector *et al.*, 2001; Davis, 2009).

Ecosystem functioning is driven, beyond the sheer number of species, by community composition in terms of key functional trait (Gagic *et al.*, 2015). Communities with the same diversity, but different trait compositions, might possess different functioning characteristics. Communities probably harbor very different traits depending on whether they are recent assemblages drawn from the regional pool, or if species have adapted to the local environment and to the other species, through various mechanisms including plasticity, niche-construction and evolution (Kylafis & Loreau, 2011; Hendry, 2016; Meilhac *et al.*, 2020). In particular, evolutionary changes may be important on ecological timescales (Davis *et al.*, 2005; Hendry, 2016), and there is mounting evidence that species can adapt rapidly to environmental changes and to the presence of competitors or predators (Thompson, 1998; Faillace & Morin, 2016; Kleynhans *et al.*, 2016; Hart *et al.*, 2019; Meilhac *et al.*, 2020). By altering species trait composition, such community adaptation may impact the existence, magnitude and shape of BEF relationships.

Even though BEF studies are traditionally conducted from an ecological perspective, long term grassland experiments (Reich *et al.*, 2012; Meyer *et al.*, 2016) and microbe experiments (Bell *et al.*, 2005) found that biodiversity-yield relationships change through time. The most recent studies have explicitly highlighted a role of evolution in modifying biodiversity-yield relationships: in grasslands (Zuppinger-Dingley *et al.*, 2014; van Moorsel *et al.*, 2018) and with microbes, using experimental evolution (Fiegna *et al.*, 2014, 2015). However, results have proven quite variable, prompting a plea for more theoretical investigations (Fiegna *et al.*, 2015).

Here we propose a theoretical evaluation of the consequences community adaptation may have for BEF relationships. We use a general modeling approach to address the three types of BEF relationships highlighted above (biodiversity-productivity, biodiversity-stability, and biodiversity-invasion). We compare two contrasted types of communities: (i) random communities, in which only ecological processes control species trait composition, with no evolutionary dynamics, and (ii) co-adapted communities, in which species traits composition have further been shaped by micro-evolution (species adaptation to the environment and to other species). Specifically, species have adjusted their traits and are at (co)evolutionary equilibrium (Christiansen, 1991). Real-life communities would harbour various degrees of co-adaptation in between these two limiting cases. Recently-founded or perturbed ecosystems, such as artificially assembled communities, are probably closer to the random case. In contrast, ecosystems that have long remained in relatively constant conditions, such as primary old-growth forests, may be closer to co-adapted communities.

As species coexistence and eco-evolutionary dynamics depend on the type of ecological interactions (Mouquet *et al.*, 2002; Chave *et al.*, 2002; Calcagno *et al.*, 2017), we model communities governed by four contrasted scenarios of ecological interactions, representative of classical coexistence mechanisms (Doebeli & Dieckmann, 2000; Calcagno *et al.*, 2017): two scenarios based on resource competition (one symmetric, one asymmetric), one life-history trade-off scenario, and a trophicchain scenario. In each case, several functioning metrics are computed to evaluate BEF relationships. This general approach allows to evaluate the extent to which the consequences of community adaptation are general or depend on particular types of metrics and ecological interactions.

We report clear influences of community adaptation on each of the three BEF relationships investigated, highlighting how co-adaptation impacts species trait distribution and, in turn, functioning properties. Although conclusions may depend importantly on the type of ecological interaction scenario considered, general predictions regarding the consequences of community adaptation are formulated, and discussed in light of available experimental evidence.

## Material and methods

### Ecological scenarios and traits

In natural ecosystems species are engaged in various interactions, within the same trophic level (horizontal interactions) and among different trophic levels (vertical interactions), at different spatio-temporal scales. The dominant form of species interaction may differ across communities (Chave *et al.*, 2002), and some studies have argued that generalist predation, exploitative competition and simple three-species food chains compose the common backbone of interaction networks (Mora *et al.*, 2018). To reflect this diversity, we here considered a set of four contrasted ecological scenarios (Fig. 1a), based on classical species coexistence models, and spanning the range from completely horizontal symmetric interactions to completely vertical asymmetric interactions. The first two scenarios describe competition for resources. The *Niche* scenario is a classical model of symmetric competition along an axis of niche differentiation (Doebeli & Dieckmann, 2000; Calcagno *et al.*, 2017). The *Bodysize* scenario introduces interference and asymmetric competition, based on *e.g.* size differences (Rummel & Roughgarden, 1985; Doebeli & Dieckmann, 2000). The third scenario (*LH-tradeoff*) models a life-history trade-off, describing the strongly asymmetric competition between species good at colonizing empty habitat and species locally dominant, along a competitive hierarchy (Tilman, 1994; Calcagno *et al.*, 2006, 2017). Last, the fourth (*Trophic*) scenario describes a size-structured trophic chain, based on the model introduced by Loeuille & Loreau (2005).

**Fig. 1.**
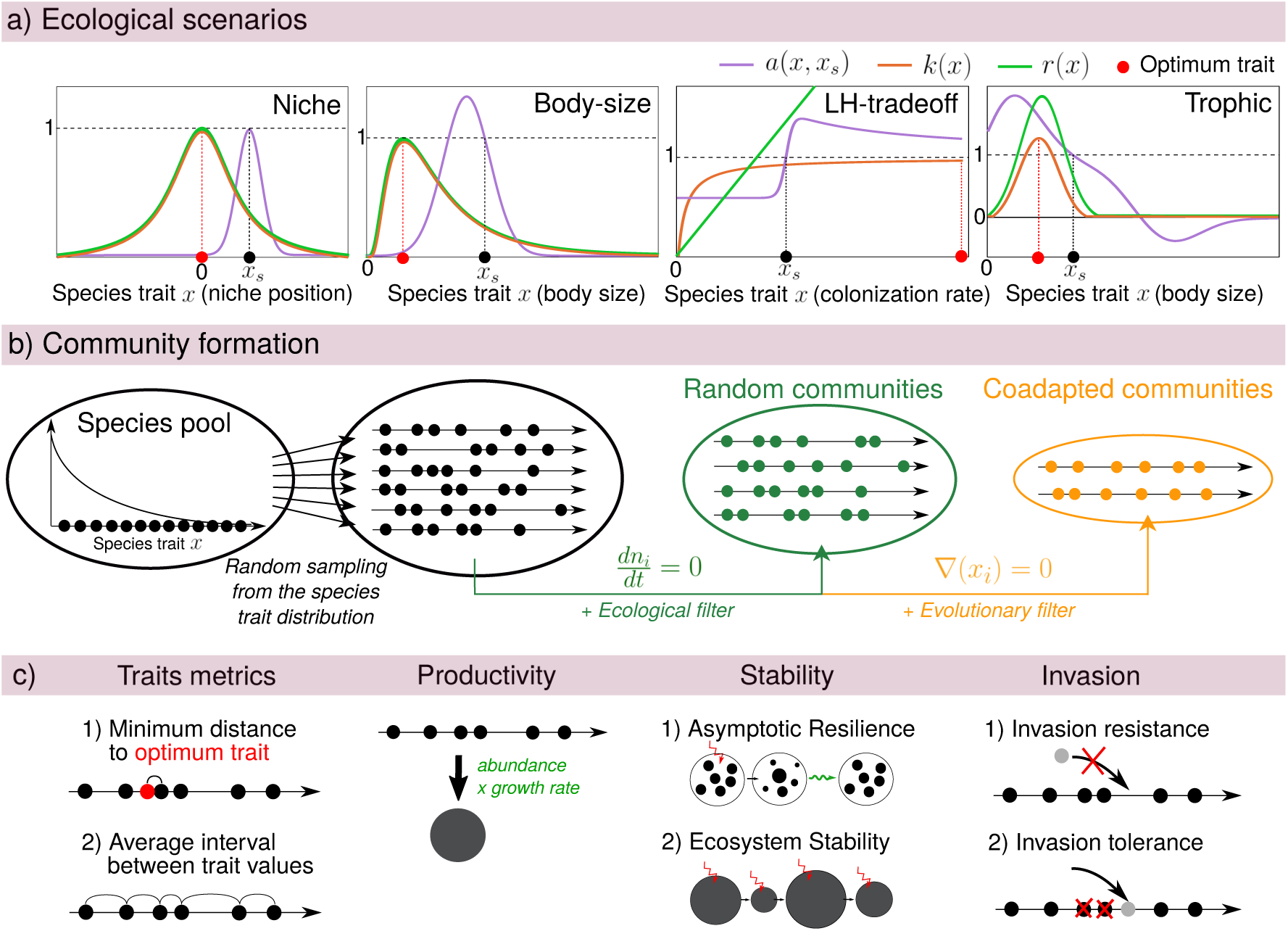
(a) Ecological parameters for the four scenarios considered: intrinsic growth rate *r*(*x*) (green), mono-specific abundance *k*(*x*) (orange) and competitive impact *a*(*x, x_s_*) (purple) of species *x_s_* over species *x* as a function of species trait *x* for the four scenarios. By definition the intra-specific competition is *a*(*x_s_, x_s_*) = 1. Red dots indicate the optimum ecological trait, *i.e.* trait that maximizes mono-specific abundances. (b) Community formation. Species are sampled from a species pool within a given distribution. An ecological filter is then applied so that only the ecologically existing communities are kept (with no null abundance), and form the random communities. Then, species evolution towards their evolutionary equilibrium filters out some species, leading to co-adapted communities. (c) For each community (random or co-adapted), we measure two species trait metrics (minimum distance to optimum trait, and average interval between trait values) and the three types of functioning properties: (i) productivity measured by species abundances time species growth rates, (ii) stability, with asymptotic resilience (return rate to equilibrium) and ecosystem stability (reflecting changes in abundances over time), and (iii) response to invasion, with invasion resistance (probability of non-establishment of a foreign species) and tolerance to invasion (probability of non resident extinction following an invasion).

In each interaction scenario, species are characterized by one key trait, denoted *x* (Fig. 1a). In the *Niche* scenario, the trait represents niche position along the continuum of resources, and interspecific competition thus decreases with trait difference (niche differentiation). In the *Body-size* scenario, the trait is body size: species with similar size compete more intensely, and bigger species have a competitive advantage over smaller ones. In the *LH-tradeoff* scenario, species trait is the colonization rate: species with greater trait value are more apt at colonizing empty patches, but also more susceptible to be competitively displaced from occupied patches (Calcagno *et al.*, 2006). Last, in the *Trophic* scenario, species trait is body mass: body mass influences growth and metabolic rates, and species preferentially consume species that are smaller, with some optimal mass difference (Loeuille & Loreau, 2005).

After appropriate reformulations (Supporting information (S.I.) Section 1), all models can be set in the common Lotka-Volterra form:

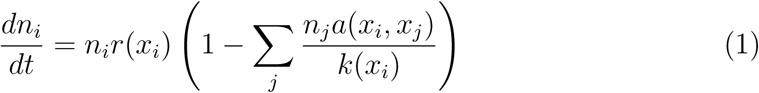

with *n_i_* the abundance of species *i*, that denotes, depending on scenario, either a number of individuals (*Niche*), a biomass (*Body-size* and *Trophic*) or a fraction of occupied sites (*LH-tradeoff*).

The three functions included in equation (1) allow to describe species demography and inter-specific interactions: *r*(*x_i_*) is the intrinsic growth rate of species *i* that governs the ecological timescale; *a*(*x_i_*, *x_j_*) is the impact that a variation in species *j* abundance has on the per capita growth rate of species *i*, normalized by the intra-specific interaction (see S.I., Section 1.1); and *k*(*x_i_*), usually called the carrying capacity, quantifies the resistance to density dependence of species *i*. In all scenarios but the *Trophic* one, it is also the equilibrium abundance reached by the species if growing alone in the community, or in other words the mono-culture abundance (Loreau & Hector, 2001). The shape of the functions for each scenario differ in important ways, as represented in Fig. 1a. See S.I. Section 1 for a complete description of each scenario.

Evolution would often favor certain trait values that are better adapted to the current habitat; this is described by the mono-specific abundance function *k*(*x*), which defines the optimum trait value, as represented by the red dots in Fig. 1a. The relationship between trait value and mono-specific abundance may have an intermediate optimum (*Niche, Body-size* and *Trophic*) or be open-ended (*LH-tradeoff*), see red dots positions in Fig.1a. Sometimes, inter-individual interactions and competition may counteract evolution towards optimal trait values, in particular in the *LH-tradeoff* scenario, in which evolution effectively results in traits with comparatively low mono-specific abundances (Calcagno *et al.*, 2017).

Species traits, through functions *r*, *k* and *a*, determine species interactions and overall ecosystem and evolutionary dynamics. Note that we consider here species that coexist stably and have distinct ecological traits. For each scenario, one or two parameters controlling the shape of the functions were systematically varied to ensure that conclusions were robust to parameter changes (all details and the parameter ranges explored are provided in S.I., Section 2).

In the *Niche* scenario, we varied the width of the competition function (Fig. 1a) keeping the width of the mono-specific abundance function constant (Doebeli & Dieckmann, 2000). In the *Body-size* scenario, we varied both the width of the competition function and its skew (level of competitive asymmetry; see Rummel & Roughgarden, 1985; Doebeli & Dieckmann, 2000). In the *LH-tradeoff* scenario, we varied the intensity of the tradeoff, and the level of competitive preemption (Calcagno *et al.*, 2006). Finally, in the *Trophic* scenario, we varied the level of interference competition and the width of the consumption function (Loeuille & Loreau, 2005). In the figures, for clarity, only three contrasted and representative parameter sets are presented per scenario.

### Random and co-adapted communities

The process of community formation is sketched Fig. 1b. For diversity levels (*N*) between 1 and 10, sets of species were drawn randomly from a regional pool. The ecological equilibrium with *N* species was computed from equation (1), and the community was retained if all species persisted at equilibrium (see S.I. Section 3 for details). This was repeated until obtaining, for each diversity level, 1,000 such random communities. The distribution of species trait values in the regional pool was chosen to minimize information content (maximum entropy; Jaynes, 1957), while being representative of typical trait values expected for the corresponding ecological scenario and parameter set. This is a generic approach but, of course, there are many ways in which diversity gradients can be generated in nature and experiments. We tried alternative methods to assemble random communities, and conclusions were little affected (see S.I. Section 3). For some scenarios and parameter sets, no feasible community could be found beyond some diversity level, in which case we stopped at the highest feasible level.

Whereas random communities are only constrained by ecological processes (regional pool and local competitive exclusion), co-adapted communities met the additional constraint that species traits are at (co)evolutionary equilibrium (“evolutionary filter”; Fig. 1b). We computed, for each species in a community, the selection gradient (Christiansen, 1991), defined as ds

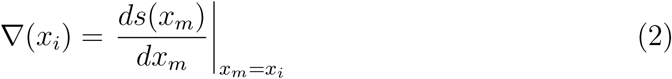

where *s*(*x_m_*) is the fitness (growth-rate) of a rare phenotype *x_m_*. Note that fitness is density- and frequency-dependent and varies with community composition (species trait and abundances).

If ∇(*x_i_*) *>* 0, selection acts to increase the trait value, whereas if ∇(*x_i_*) *<* 0 smaller values are selected for. When all selection gradients are simultaneously cancelled in a community, species have attained an evolutionary attractor and are at equilibrium with respect to first-order selection (Christiansen, 1991). This approach assumes heritable trait variation and sufficiently small phenotypic variance within each species. We thus generated, for each scenario, parameter set and diversity level, all possible co-adapted communities (most often, only one), in the sense of eq. (1) and (2). See S.I. Section 3 for detailed methods.

### Biodiversity-functioning relationships

For each generated community, we computed several properties of interest (Fig. 1c; full list in S.I. Section 4) to investigate the three BEF relationships. We present results based on the properties describe herefter, as conclusions were similar based on the others. First, community productivity (Tilman, 1999; Loreau & Hector, 2001) was measured as the species average rate of production (positive contribution to growth rate) in the community. Second, ecological stability was assessed in two ways. We computed the classical asymptotic resilience (May, 1973; Arnoldi *et al.*, 2016), *i.e.* asymptotic rate of return to equilibrium of the community after a perturbation, and the community stability (May, 1973; Ives *et al.*, 1999), *i.e.* the inverse of the coefficient of variation of total community abundance under sustained environmental noise. Finally, to study the response to invasions, we also used two properties. The first is the resistance to invasion (Elton, 1958; Hector *et al.*, 2001), *i.e.* the probability that a random alien species, introduced at low abundance, fails to establish in the community. The second was the tolerance to invasion (Elton, 1958), *i.e.* the proportion of species that, following a successful invasion, were not driven to extinction. Details on the mathematical computation of each metric are presented in S.I. Section 4.

For each metric, diversity level, scenario and parameter set, we computed the average value over the 1,000 random communities, and over the few (or, most often, the unique) co-adapted communities. To ensure that average differences represented large effect sizes, we further computed the percentile, in the distribution of values over random communities, corresponding to the value of co-adapted communities. Our results showed that co-adapted communities often lie in the tail of the distribution of random communities, for all metrics (see S.I. Section 5). Average differences between co-adapted and random communities were thus large relative to the variability of random communities. For this reason we only present average values in the Figures.

The above metrics were correlated to species richness to produce BEF relationships and compare random and co-adapted communities. Since the impacts of co-adaptation are mediated by changes in trait values, we compared the structure of co-adapted and random communities. We then computed the average absolute difference in trait space between the two, as a measure of the strength of the evolutionary filter. We summarized trait compositions using two additional quantities (Fig. 1c). The first was the minimum distance to the optimal trait value (red dots in Fig. 1a), that reflects how well the better performing species is adapted to the habitat. The second was the average trait interval between species (trait range divided by number of species minus one), that indicates how “packed” species are in trait space. More details are provided in S.I. Section 4.

## Results

### Biodiversity-Productivity

Random and co-adapted communities differed in productivity at low diversity levels, but at higher diversity levels, differences were more modest (Fig. 2). Moreover, in all scenarios except the *LH-tradeoff*, the effect of community adaptation was to increase productivity. Those two observations explain the quantitative differences in biodiversity-productivity relationships between random and co-adapted communities.

**Fig. 2.**
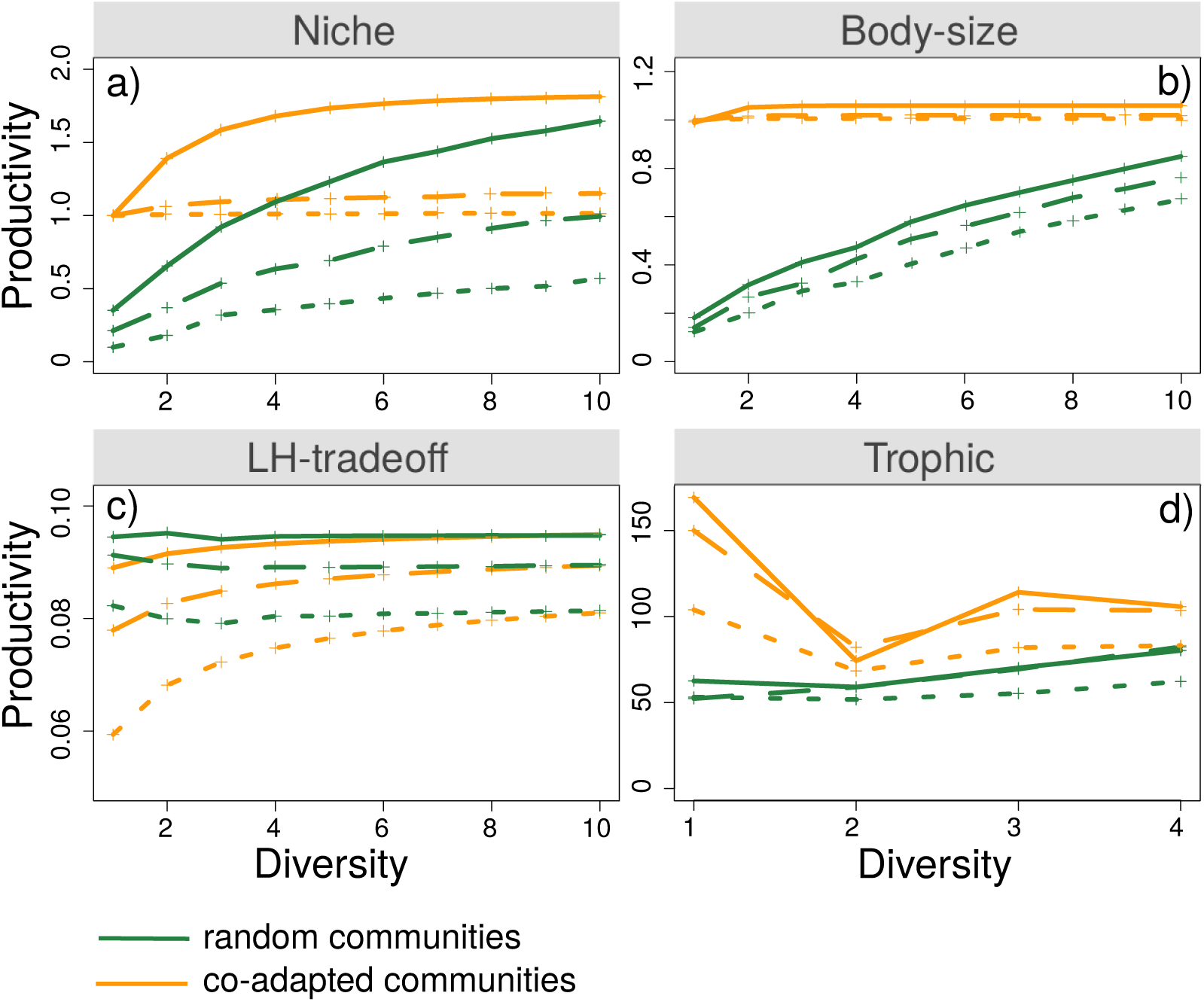
Productivity as a function of diversity under the the four scenarios and the two community adaptation levels. Three sets of parameters are used for each scenario, represented by the three different line types. Parameter values are given in S.I. Section 2 together with other explored parameter sets (not shown for the sake of clarity). Each point represents an average over 1000 random communities or the only or few co-adapted communities.

Qualitatively, co-adaptation affected the biodiversity-productivity relationship in all four scenarios (Fig. 2). The impact could be as spectacular as a slope inversion. For instance, the *LH-tradeoff* scenario, unlike the other scenarios, generated mildly negative biodiversity-productivity relationships in random communities (see also Loreau, 2010), while in co-adapted communities, they switched to markedly positive for all parameter sets (Fig. 2c). Conversely, the *Trophic* scenario generated a classical positive biodiversity-productivity relationship in random communities, but the relationships switched to negative in co-adapted communities (with oscillations between odd and even diversity levels, caused by trophic cascades; Fig. 2d). The possibility of such inversions of biodiversity-production relationships has, to the best of our knowledge, never been reported so far.

In the remaining scenarios, those based on resource competition, biodiversity-productivity relationships were always positive – at least slightly – irrespective of co-adaptation (Fig. 2a,b). However, the shape of the relationships differed markedly between random and co-adapted communities: the increase in productivity with diversity was close to linear or gradually slowed down with diversity, whereas in co-adapted communities, the relationships saturated very quickly, reaching almost maximum productivity at low diversity levels and then plateauing, especially in the *Body-size* scenario (Fig. 2b).

### Species trait composition

As shown in Fig. 3, random and co-adapted communities exhibited systematic differences in their trait composition and structure. The specifics differed across ecological scenarios, but general trends can be identified. First, random communities are generally less packed than co-adapted ones, as can be seen by the slopes lower than one in Fig. 3), indicative of broader trait ranges in random communities. Second, the difference was maximal at low diversity and tended to vanish as diversity increases. Increasing diversity made random and co-adapted communities converge to similar trait distributions on average (aligning on line *x* = *y* in Fig. 3), with one exception in the *Trophic* scenario. As a result, the impact of community adaptation on community structure, i.e. the strength of the evolutionary filter, globally declined with the number of species (insert panels in Fig. 3).

**Fig. 3.**
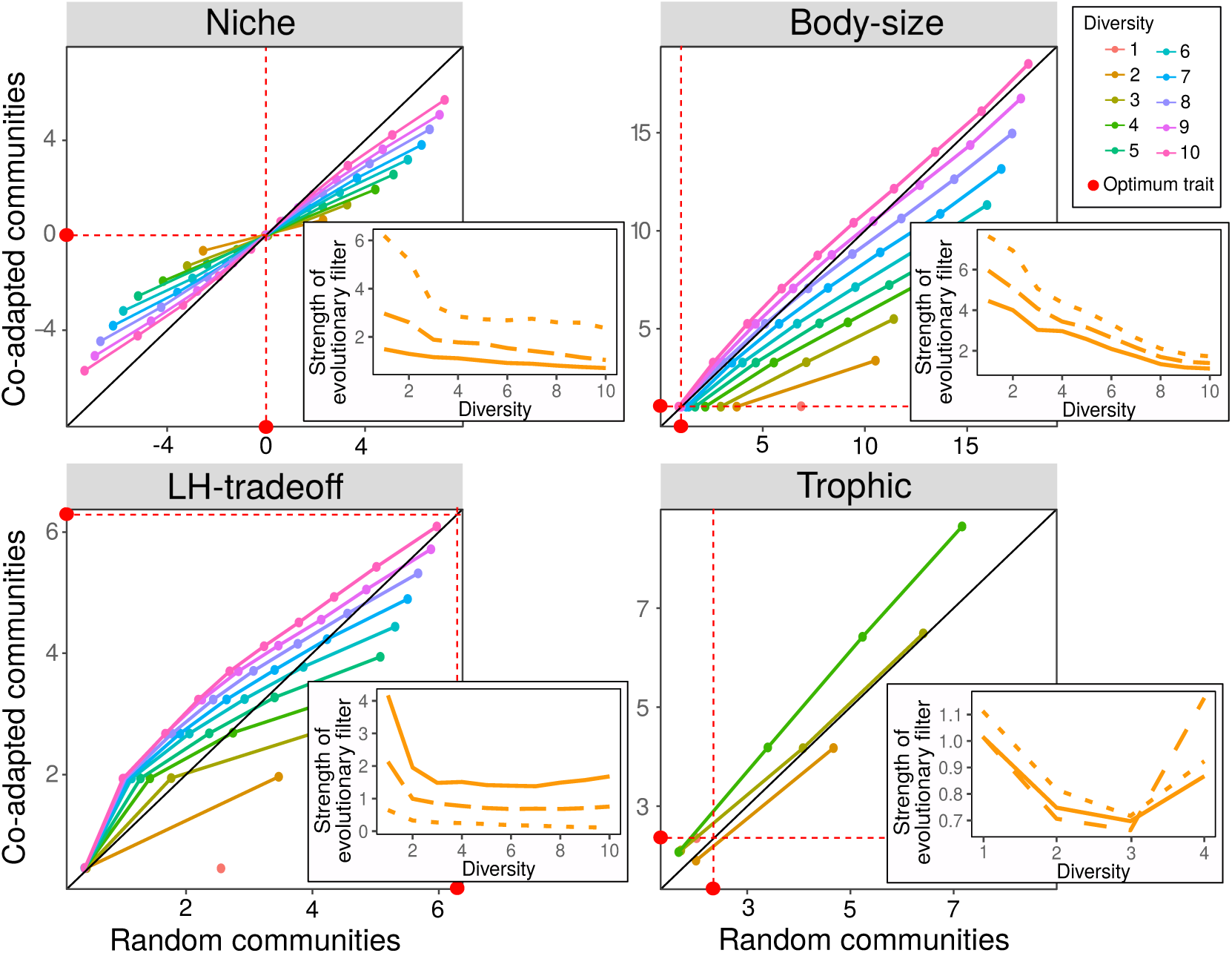
Comparative structures of random and co-adapted communities, in the four ecological scenarios. For each diversity level, species were ranked by trait value, and trait values at each species rank were averaged over all communities. The average trait values at each species rank in co-adapted communities was plotted against the corresponding values in random communities, and connected with colored lines for each diversity level (see legend). As a consequence, lines close to the first diagonal indicate very similar trait compositions in the two community types. Slopes smaller than one indicate greater trait dispersion in random than in co-adapted communities, while slopes greater than one indicate the opposite. Only one parameter set was showed in each scenario (the one corresponding to the wide dotted lines in other figures), for clarity, as patterns are similar in other parameter sets. Optimal trait values (see Fig. 1) are also shown on both axes (red dots). In inserts, we represented the strength of the evolutionary filter as a function of diversity, for all three parameter sets (each with a different line type). This was computed as the absolute trait difference between random and co-adapted communities, per species rank, averaged over all ranks and all communities.

More specifically, in random communities, the chance to find a highly performing species inevitably increased with the number of species, so that the minimum trait distance to the optimum decreased with diversity in all scenarios (Fig. 4a-d). Concomitantly, the average distance between species decreased sharply with species richness (Fig. 4e-h), reflecting the greater species packing. In contrast, a close-to-optimal species was always present in co-adapted communities (constant minimum distance to optimum trait Fig. 4a,b,d), except for the *LH-tradeoff* scenario in which evolution did not drive species to the optimum trait (Fig. 4c). Community adaptation also made the level of species packing virtually constant irrespective of species number (Fig. 4e-h; see also Fig. 3).

**Fig. 4.**
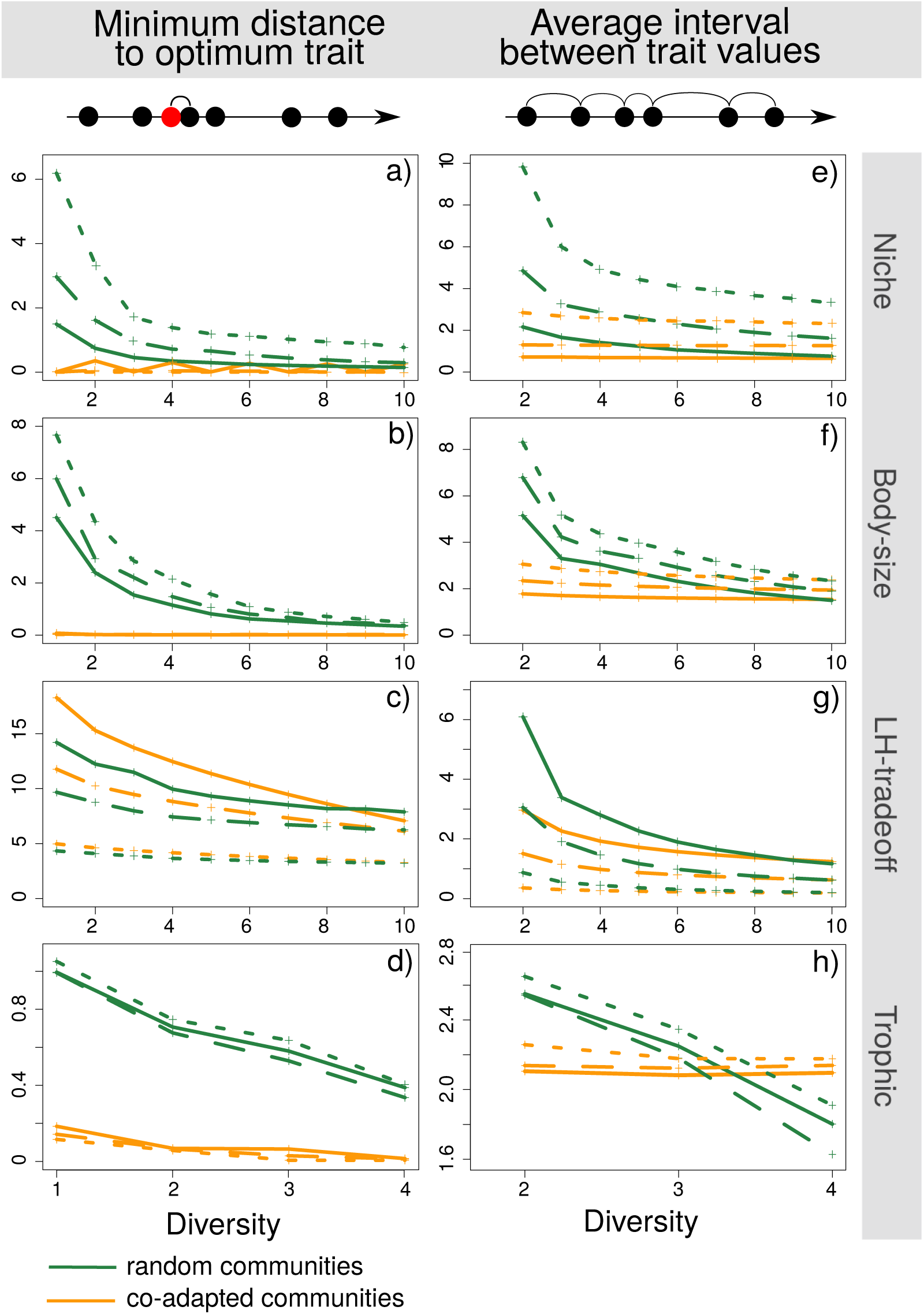
Minimum distance to optimum trait (a-d) and average interval between trait values (e-h) as a function of diversity under the four scenarios and the two types of communities. Three sets of parameters are used for each scenario, represented by the three different line types. Parameter values are given in S.I. Section 2 together with other explored parameter sets (not shown for the sake of clarity). Each point represents an average over 1000 random communities or the only or few co-adapted communities.

### Biodiversity-Stability

Asymptotic resilience, in all four ecological scenarios, declined with diversity (Fig. 5a-d). Moreover, the biodiversity-stability relationships were similarly negative, regardless of community adaptation, even though co-adapted communities were generally more stable than random ones.

**Fig. 5.**
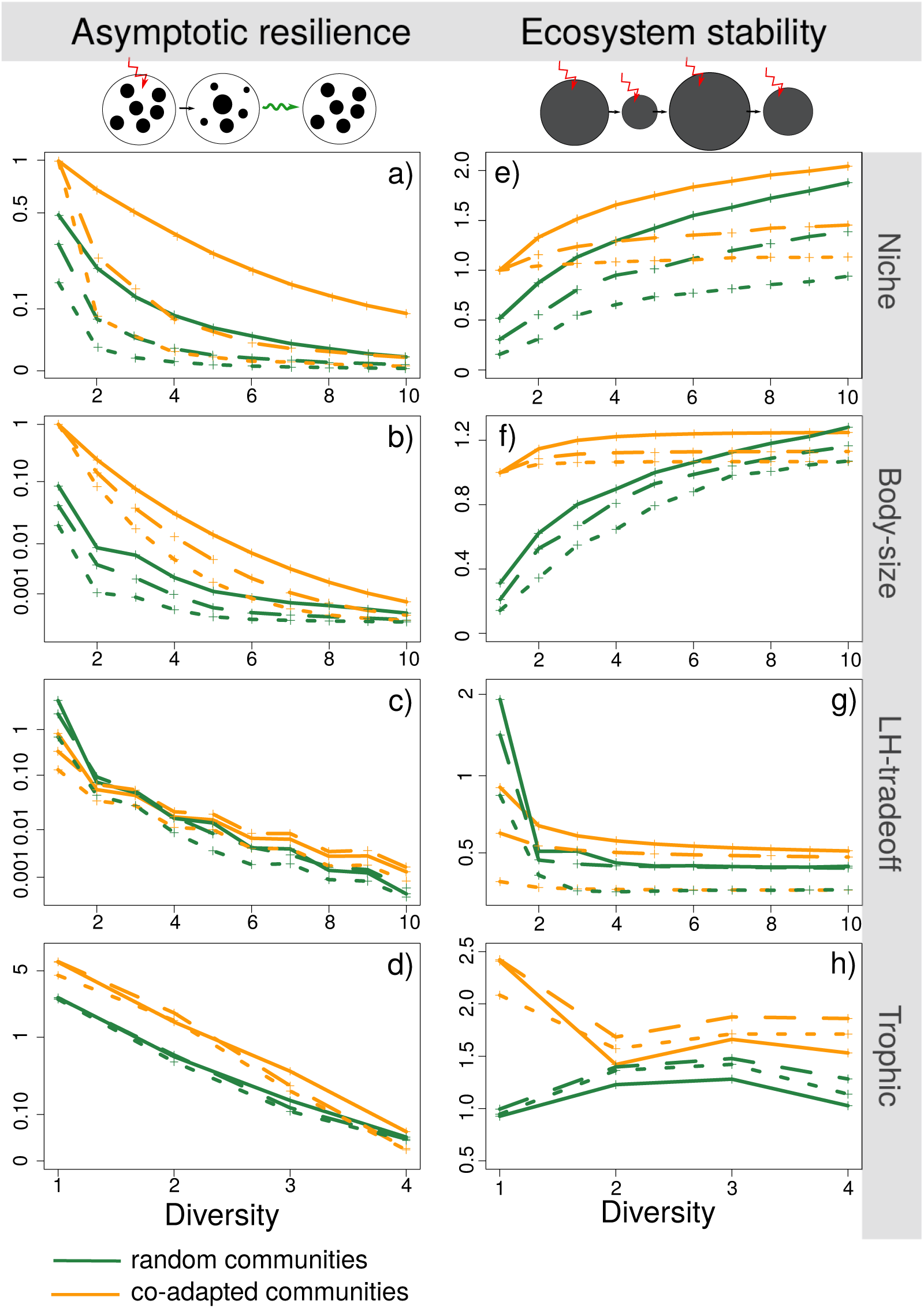
Asymptotic resilience (a-d) and ecosystem stability (e-h) as a function of diversity under the four scenarios and the two community adaptation levels. Asymptotic resilience is represented in log-scale, which does not modify the inter-pretation on co-adaptation effect and allows to better visualize the consequences of co-adaptation. Three sets of parameters are used for each scenario, represented by the three different line types. Parameter values are given in S.I. Section 2 together with other explored parameter sets (not shown for the sake of clarity). Each point represents an average over 1000 random communities or the only or few co-adapted communities.

Ecosystem stability, was also higher overall in co-adapted than in random communities (Fig. 5e-h). However, unlike asymptotic resilience, it had different responses to diversity depending on ecological scenario. It increased with species richness in the two scenarios based on resource competition (*Niche* and *Body-size*), but decreased with species richness in the *LH-tradeoff* and *Trophic* scenarios. In all cases, unlike asymptotic resilience, the variation of ecosystem stability with species richness was strongly affected by co-adaptation, and the patterns were quite consistent with those observed for total productivity (Fig. 2), except for the *LH-tradeoff*.

### Biodiversity-Invasion

Resistance to invasion (Fig. 6a-d) presented consistent trends in the four ecological scenarios. First, it increased with species richness, reflecting classical expectations. Second, co-adapted communities were generally more resistant to invasion than random ones, at any species richness level, reflecting the concentration of species around trait optima, which leaves only more peripheral niches available for potential invasive species. This difference was also quite in line with common expectations, but it could vanish, or even reverse for some parameter values, in the *LH-tradeoff* scenario (Fig. 6c). Overall, the biodiversity-invasion relationships were thus similar regardless of co-adaptation.

**Fig. 6.**
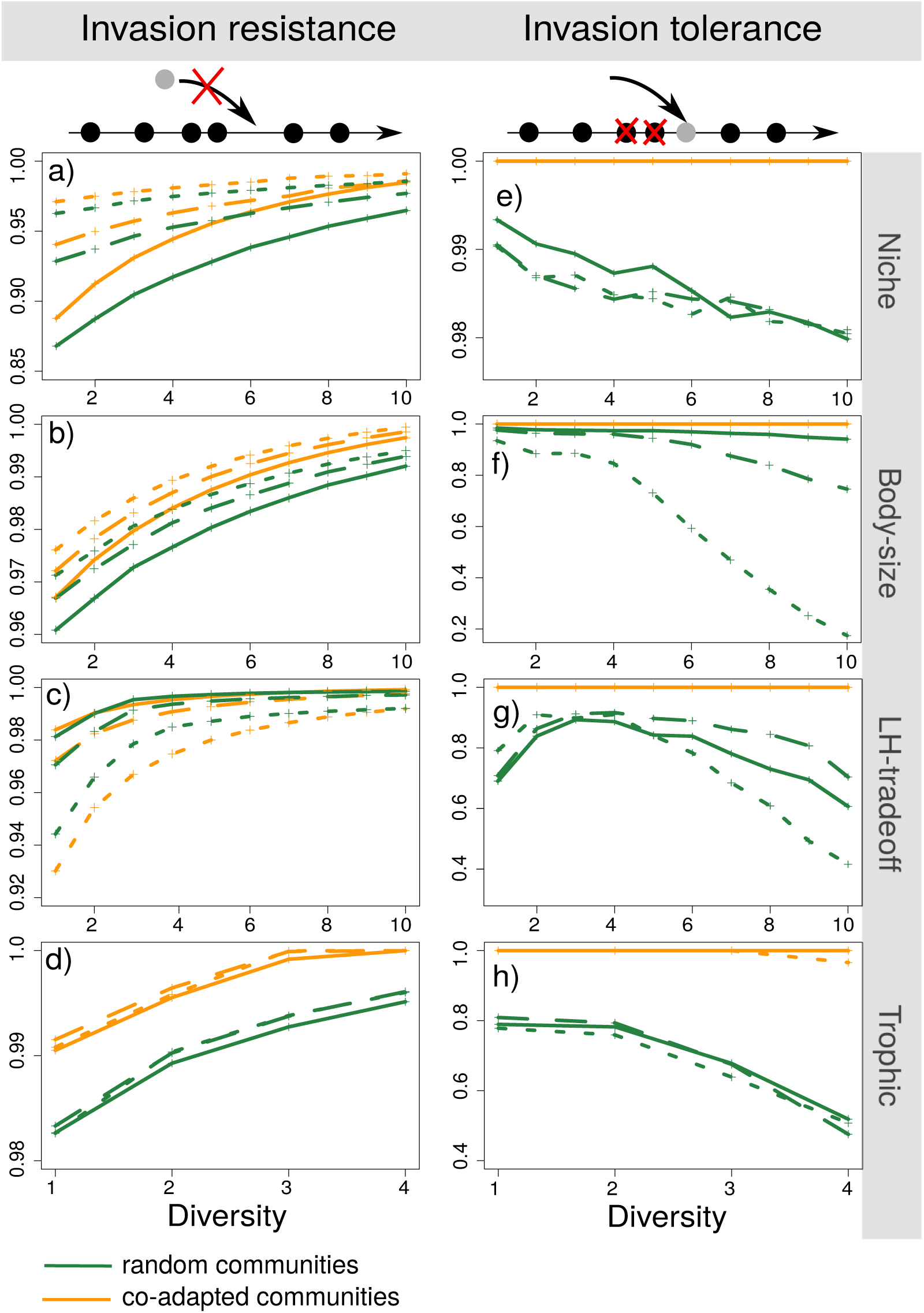
Probability that a foreign species does not get installed into a community, i.e. resistance to invasion (a-e), and proportion of resident species that do not undergo a secondary extinction, i.e. tolerance to invasion (f-j), as a function of diversity, under the four scenarios and the two community adaptation levels. Three sets of parameters are used for each scenario, represented by the three different line types. Parameter values are given in S.I. Section 2 together with other explored parameter sets (not shown for the sake of clarity). Each point represents an average over 1000 random communities or the only or few co-adapted communities.

The effects of co-adaptation were much more dramatic and consistent when looking at tolerance to invasion (Fig. 6e-g), with a pronounced interaction between the effects of diversity and community adaptation. In random communities tolerance to invasion steeply declined with species richness, meaning that successful invasions were more and more harmful (in terms of diversity loss) as diversity increased (Fig. 6e-g). In contrast, co-adapted communities were in all cases more tolerant to invasion than random communities: they retained their almost perfect tolerance to invasion as diversity increases, the biodiversity-invasion relationship being essentially flat.

## Discussion

Natural and anthropized ecosystems present tremendous variation in their diversity and composition (species richness and trait values), and also in the degree to which component species are adapted to the local environmental conditions and to one another, here referred to as community adaptation. While it is clear that diversity is an important determinant of ecosystem functioning, we still know little about how the level of community adaptation might impact BEF relationships (Fiegna *et al.*, 2015; Hendry, 2016). In this study we addressed this question with a general modelling approach, systematically comparing random and co-adapted communities with respect to three BEF relationships (biodiversity-productivity, biodiversity-stability and biodiversity-invasion) and across four classical scenarios of ecological interactions.

We found that community adaptation had an impact on all BEF relationships, but that the nature and extent of the impact depend on both the metrics and the scenarios considered for species interactions. Overall, the biodiversity-productivity and biodiversity-invasion relationships were strongly impacted by community adaptation, while the biodiversity-stability relationships were much less so. Indeed, co-adapted communities, at any species richness, tended to be more dynamically stable in terms of asymptotic resilience than random ones, but there was little interaction with diversity: BEF relationships looked qualitatively very similar in random and co-adapted communities. This suggests that the connection between diversity and dynamical stability is a rather universal property in such systems of interacting species, largely insensitive to the details of species interactions and to community adaptation. Co-adapted communities were more dynamically stable, which likely reflects the fact that co-adaptation brings traits closer to optimal values (Fig. 4a-d), entailing faster returns to equilibrium (see also Loeuille, 2010). Consistent with this interpretation, the only case where co-adapted communities were on average less dynamically stable than random ones occurred at low diversity in the *LH-tradeoff* (Fig. 5c), a case where co-adaptation pushes traits away from the optimum (Fig. 1a and 3). When looking at ecosystem stability (May, 1973; Ives *et al.*, 1999; Arnoldi *et al.*, 2016), BEF relationships differed more importantly between co-adapted and random communities, but these differences closely mirrored those observed for biodiversity-productivity relationships (Fig. 5e-h). This suggests that variation in ecosystem stability were linked to variation in total productivity and the total biomass of species, but, beyond that, were little impacted by community adaptation in a direct manner (see Ives *et al.*, 1999), especially for the *Niche*, *Bodysize* and *Trophic* scenarios. Consistent with this interpretation, overall Spearman correlation coefficients between productivity and ecosystem stability were, across all communities, 0.97, 0.97, 0.65 and 0.79 for the *Niche*, *Body-size*, *LH-tradeoff*, *Trophic* scenarios, respectively.

Biodiversity-productivity relationships were both quite variable across ecological scenarios and strongly impacted by community adaptation. The impact could be as pronounced as a slope inversion between random and co-adapted communities, with switches from positive to negative (*Trophic* scenario; Fig. 2d) or from negative to positive (*LH-tradeoff* scenario; Fig. 2c). In all cases, changes in the shape of biodiversity-productivity relationships were mostly driven by low diversity levels, at which co-adapted communities differed importantly for random ones, while at higher diversity levels community adaptation had modest impact on average (Fig. 2). Theoretical (Mazancourt *et al.*, 2008) and experimental (terHorst *et al.*, 2018; Scheuerl *et al.*, 2020) studies have found that biodiversity inhibits the evolution of species traits (but see Jousset *et al.* (2016), and our *Trophic* scenario; Fig. 3d). This can be attributed to the strong constraints on species trait distributions in rich communities, due to species interactions and persistence conditions (ecological filter), which leaves adaptive evolution much less room to alter species traits. As a result, co-adapted and random communities are more and more similar as the number of species increases, i.e. the strength of the evolutionary filter decreases (see Fig. 3: its presence makes less and less of a difference for ecosystem functioning.

In most ecological scenarios (*Niche*, *Body-size* and *Trophic*), co-adapted communities at low diversity levels were more productive than random ones on average (Fig. 2a,b,d). Therefore, community adaptation weakened the positive biodiversity-productivity relationships observed in random communities, making those shallower, or even reversing them to negative (*Trophic*). In the *LH-tradeoff* scenario, low diversity co-adapted communities were on the contrary less productive than random ones, so that the effect of community adaptation was opposite: co-adapted communities exhibited a positive biodiversity-functioning relationships, whereas a slightly negative relationship is predicted in random communities (see also Loreau, 2010).

The qualitative effect of community adaptation on biodiversity-productivity relationships could thus be determined, to a large extent, from the overall direction of selection in isolated species (monocultures), either towards higher productivity (*Niche*, *Body-size*, *Trophic* scenarios), weakening positive relationships or switching them to negative, or towards lower productivity (*LH-tradeoff* scenario), switching negative relationships to positive.

Community adaptation also affected the mechanisms underlying biodiversity-productivity relationships. In the two scenarios describing competition for resources (*Niche* and *Body-size*) and in the *Trophic* scenario, low diversity co-adapted communities were more productive because adaptive evolution favoured traits that were close to the optimum (Fig. 4a,b,d). At higher diversity levels, more and more species were kept farther away from the optimal trait value, and thus intrinsically less productive. Such a change in the intrinsic productivity of species as diversity increases is often called a “selection effect” (Loreau & Hector, 2001), but here, following Zuppinger-Dingley *et al.* (2014), we will call it a “sampling effect” to avoid confusion. In random communities, this sampling effect was positive, as usually expected, and contributed importantly to the positive biodiversity-productivity relationship: richer communities were more likely, by chance, to harbor species that were intrinsically more productive in the habitat (Fig. 4a,b,d, blue curves). In co-adapted communities, the sampling effect was much reduced or entirely absent, owing to the effect of co-adaptation explained above. In contrast, the average trait distance between adjacent species (the level of niche packing) declined sharply with diversity in random communities, but much less so in co-adapted communities (Fig. 4e-h). Therefore, the level of species trait complementarity was less sensitive to diversity with community adaptation. Altogether, this suggests positive biodiversity-productivity relationships should be more driven by complementarity effects, and less by sampling effects, in co-adapted communities compared to random communities.

There is good experimental evidence that biodiversity-productivity relationships do change over time. In grassland experiments, the positive biodiversity-productivity relationships were reported to become steeper (Reich *et al.*, 2012), or sometimes flatter (Meyer *et al.*, 2016, for most of its biomass-related metrics), over several years. A study of decomposing microbial communities observed a decline in productivity and a flattening of the biodiversity-productivity relationship over several days (Bell *et al.*, 2005). Several of these studies (Bell *et al.*, 2005; Reich *et al.*, 2012; Meyer *et al.*, 2016) deal with short timescales and are not directly relevant to address community adaptation, since observed changes are generally attributed to transient ecological mechanisms, such as below-ground feedbacks or resource depletion. Direct comparison with our results, in which transient dynamics have been sorted out, is thus difficult. Fortunately, more recent analyses of the longest grassland experiments have looked for, and found, evidence of character displacement and evolution of niche differentiation, even on relatively short timescales (Zuppinger-Dingley *et al.*, 2014; van Moorsel *et al.*, 2018). This suggests that the evolutionary effects analyzed in this work might have begun to play a role. Interestingly, it was observed that biodiversity-productivity relationships assembled from co-selected species were higher at low diversity, but saturated faster with diversity, thus being more concave (van Moorsel *et al.*, 2018). This is strikingly reminiscent of our predictions in the standard “resource competition” scenarios (*Niche* and *Body-size*; see Fig. 2a,b). Furthermore, it was found that species evolving in mixed assemblages (thus approaching a state of community adaptation) elicited more complementarity effects, and less sampling effects, than assemblages of non-co-adapted species (Zuppinger-Dingley *et al.*, 2014). This too is quite consistent with our findings. In a more direct approach, Fiegna *et al.* (2014, 2015) used experimental evolution to demonstrate that biodiversity-productivity relationships are impacted by co-adaptation in bacteria. They further showed that these changes involved an evolutionary component in species interactions, not just of individual species performances, even though the overall impact on biodiversity-productivity relationships was quite variable among experiments. These studies thus clearly support a role for community adaptation in the dynamics of BEF relationships as highlighted here.

Regarding the biodiversity-invasion relationships (Fig. 6), the effects of community adaptation were much more consistent across ecological scenarios than for biodiversity-productivity relationships. Resistance to invasion increased, and tolerance to invasion decreased, with diversity under all scenarios, which conforms well to general expectations (Levine, 2000; Hector *et al.*, 2001; Davis, 2009). The impact of community adaptation was moderate for resistance to invasion (probability of establishment) but spectacular for tolerance to invasion (number of secondary extinctions) (Fig. 6e-h), highlighting that different invasion properties can behave quite differently. Indeed, resistance to invasion was only slightly impacted by community adaptation, the latter generally increasing invasion resistance, but with almost no interaction with diversity. In contrast, biodiversity-tolerance relationships markedly differed with community adaptation (Fig. 6e-h): while tolerance to invasion gradually decreased with species diversity in random communities, it remained virtually constant with community adaptation.

This can be understood in terms of changes in species trait distributions. In co-adapted communities, species traits were more concentrated around the optimal trait (Fig. 3 and Fig. 4e-h orange curves) and more evenly spaced (see S.I. Section 6 Fig S2), so that successful invaders tended to occupy peripheral niches at either tail of the trait distribution, which did not cause resident extinctions. However, this is not the case in random communities in which an invader might find vacant niche space anywhere along the trait spectrum, possibly very close to a resident species, thus potentially excluding the latter and triggering further secondary extinctions.

Beyond BEF relationships, these results have interesting implications for community assembly dynamics. With the possibility of species (co)evolution, invasion resistance and tolerance would both increase in between species colonization events. In other words, successful invasions would be rarer, but also more constructive in terms of community assembly: invaders would more often add to the community without driving many resident species extinct. With trait co-evolution, assembly trajectories should therefore be less eventful (fewer invasions and fewer extinctions), and more steadily progressing or “efficient”, compared to pure invasion-assembly. Although this prediction deserves further exploration, it nicely complements some earlier studies of community assembly (Rummel & Roughgarden, 1985).

Unfortunately, empirical evidence is even scarcer regarding the role of community adaptation for invasion properties than for productivity. Most studies focus on documenting the impact of invasions on the evolutionary dynamics of communities, not the other way round; yet, evolutionarily immature (e.g. insular) communities or recently assembled ecosystems such as anthropized habitats are known to be more sensitive to invasions than old species assemblages. Although this is suggestive of a protective role of co-evolution, the diversity-invasion relationship is difficult to relate to evolutionary history, as most long-co-evolved communities are highly diverse while recently assembled ones are usually species-poor (David *et al.*, 2017).

Our approach was a first attempt at combining eco-evolutionary theory ad BEF-relationships. It could be extended and improved in several ways. One important simplification was the assumption of one single trait that structures communities and is subject to evolution. It would be interesting to describe species interactions as governed by multiple traits, which is probably more realistic and might result in more complex eco-evolutionary dynamics (Vasconcelos & Rueffler, 2020). Similarly, we assumed no upper limit on the amount of phenotypic change a species can undergo. Trait changes between random and co-adapted communities may sometimes be quite large, especially at low diversity levels (Fig. 3). However, since genetic variation is usually not infinite, species responses to selection can be constrained. Such limits on evolution would probably weaken some of the reported effects, even though preliminary analyses suggest that results are quite robust to this (see S.I. Fig. S3 Section 7).

Overall our work highlights some potentially important consequences of evolutionary dynamics for biodiversity-functioning relationships. Through its action on species trait values, community adaptation can profoundly change the expected relationship between diversity and various ecosystem functioning properties, even in qualitative terms. This occurs because of the differential magnitude and direction of species trait evolution in poor versus rich communities, so that the ecological impact of species number interacts with the evolutionary history of communities. Therefore, BEF relationships derived from short-term experiments or observed following recent habitat perturbations might not be safely extrapolated into the future, once eco-evolutionary feed-backs have played out. This may have consequences for our understanding and prediction of the way ecosystems respond to species loss and environmental perturbations, and for ecosystem management and restoration. Eco-evolutionary theory definitely calls for more long-term and evolution-oriented studies of BEF relationships.

## Acknowledgments

FA was funded by a PhD fellowship from Université Côte d’Azur (IDEX UCA-JEDI). ML was supported by the TULIP Laboratory of Excellence (ANR-10-LABX-41) and by the BIOSTASES Advanced Grant, funded by the European Research Council under the European Union’s Horizon 2020 research and innovation program (grant agreement No 666971). PD and PJ were funded by ANR program NGB (ANR 17-CE32-0011-05). VC and PD also thank the CESAB (project COREIDS).

