## Supplementary Information for "How community adaptation affects biodiversity-ecosystem functioning relationships"

**2 Parameter sets explored and those plotted in the main article figures 6**

**3 Algorithm of community formation 7**

**4 Metrics used to quantify diversity-functioning relationships 10**

**5 Magnitude of the difference between random and co-adapted communities 15**

**6 Coefficient of variation of the average interval between trait values 17**

**7 Results with a limit on possible trait change 18**

### 1 Modelling of the four ecological scenarios

Preliminary remark: the first three ecological scenarios are models that have already been used and presented in several earlier works, such as Calcagno *et al.* (2017). More information on these can thus be found in the latter reference; only the main elements will be provided here, for easier reading. The fourth scenario is a new model derived from, and almost identical to, the size-structured trophic chain model introduced by and studied by simulation methods in Loeuille & Loreau (2005). Therefore we will provide here its mathematical derivation.

#### 1.1 Niche scenario

This scenario represents the tendency of species to exploit different parts of a resource spectrum (Dieckmann & Doebeli, 1999; Calcagno *et al.*, 2017). It models a symmetric competition along a continuum of resources, such as a range of seed sizes for granivorous birds. Trait $x$ here represents the average position of a species along that gradient, *i.e.* mean niche position (*e.g.* mean beak size). The model is usually directly formulated under a Lotka-Voltera form represented by equation (1):

$$\frac{dn_i}{dt} = r_i n_i f_i = r_i n_i \left(1 - \sum_j \frac{n_j a_{i,j}}{k_i}\right) \tag{1}$$

where $a_{i,j} = a(x_i, x_j)$ is the impact that a variation in species $j$ abundance has on the per capita growth rate of species $i$ ($\frac{1}{n_i}\frac{dn_i}{dt}$), normalized by the intra-specific interaction (so that $a_{i,i} = 1$). The closer species traits (*i.e.* the more similar resources species consume), the stronger competitive impact between them. It results that the competition is a decreasing function of the trait difference between two species, taking maximum value 1 when the two species have identical niche position. Function $a(x_i, x_j)$ is following general practice taken to be Gaussian with width $s_a$.

$$a(x_i, x_j) = \exp\left(\frac{-(x_i - x_j)^2}{s_a^2}\right) \tag{2}$$

The carrying capacity function $k_i = k(x_i)$ describes the distribution of resources available along the resource gradient, and is classically supposed to have a symmetric dome-shaped distribution centered on some optimal trait value (0 for convenience). Following Calcagno *et al.* (2017), in order to avoid some degenerate mathematical properties we use a Lorentzian function of width $s_k$

$$k(x) = \frac{1}{1 + x^2/s_k^2} \tag{3}$$

Last, $r_i = r(x_i)$ is the intrinsic growth rate of species $i$ that governs the ecological timescale, and is taken to be proportional to the mono-specific abundance $r(x) = k(x)$. The shape of the three functions presented above are represented in Fig. 1a in the main article.

Parameter $s_a$ is varied in the simulation, over the range presented in section 2, while parameter $s_k$ has been kept constant equal to 1.

##### 1.2 Body-size scenario

This second scenario (Rummel & Roughgarden, 1985) is an extension of the *niche* model in which asymmetric competition between species is used instead of symmetric competition. In this case, species trait can represents for instance species body size. It is represented by the same equation (1), with different competitive, carrying capacity and intrinsic growth rate functions. As in Calcagno *et al.* (2017), we use a log-normal carrying capacity function:

$$k(x) = \exp\left(-log(x)^2\right) \tag{4}$$

and an asymmetric Gaussian competitive function:

$$a(x_i, x_j) = \exp(d^2/s_a^2) \exp\left(\frac{-(x_i - x_j + d)^2}{s_a^2}\right) \tag{5}$$

with $s_a$ controlling the competitive function width, and $d$ the level of asymmetry. The intrinsic growth rate is also taken proportional to function $k$ : $r(x) = k(x)$. The form of those functions are represented in Fig. 1a in the main article.

Parameters $s_a$ and $d$ are varied in the simulation, over the range presented in section 2.

##### 1.3 Life History Trade-off scenario

This scenario rests on a patch-occupancy model describing the competition between species in a meta-community, arranged in a competitive hierarchy. There is a trade-off between colonization ability at the regional scale and competitive dominance at a local scale (Calcagno *et al.*, 2006, 2017).

The patch-occupancy equation given in Calcagno *et al.* (2006) can easily be rewritten under the same Lotka-Volterra form as in the two first scenarios (eq. 1) as detailed in Calcagno *et al.* (2017). The corresponding three functions $a(x_i, x_j)$, $k(x)$ and $r(x)$ are represented Fig. 1a of the main article and are defined as:

$$a(x_i, x_j) = 1 + x_j \frac{\eta(x_j - x_i)}{x_i - \eta(x_i - x_j)} \tag{6}$$

$$k(x_i) = N - \mu/x_i \tag{7}$$

$$r(x_i) = x_i N - \mu \tag{8}$$

with $\eta$ a logistic function relating the probability to win competition for a patch and the difference in colonization abilities; see Calcagno *et al.* (2017) for the full derivation.

Parameters $\alpha$ (trade-off intensity, *i.e.* the steepness of the logistic function $\eta$) and $\gamma$ (the preemption level, *i.e.* the maximum value of function $\eta$) were both varied, over the range presented in section 2. Parameters $N$ (total amount of patches in the metacommunity) and $\mu$ (patch extinction rate) were kept fixed to $N = 1$ and $\mu = 0.1$.

##### 1.4 Trophic scenario: derivation of the Lotka-Volterra form

This scenario presents a vertical trophic interaction, with species arranged in a trophic chain structured by body mass. Body mass is thus the species trait $x$. This model is taken from Loeuille & Loreau (2005) and reads:

$$\frac{1}{n_i}\frac{dn_i}{dt} = f(x_i)\sum_{j=0}\gamma(x_i - x_j)n_j - m(x_i) - \sum_{j=1}\beta(x_i - x_j)n_j - \sum_{j=1}\gamma(x_j - x_i)n_j \quad (9)$$

with $n_i$ the biomass of species $i$. Index $i = 0$ corresponds to the basal resource, and its trait $x_0$ does not evolve. The first term in the right-hand side corresponds to the consumption by species $i$ of species lower into the trophic chain, and fourth term of the consumption of species $i$ by species higher in the chain. The consumption rate function $\gamma(x_i - x_j)$ is a Gaussian of width (standard deviation) $s$, and taking maximum value when species biomasses differ by some interval $d$.

$$\gamma(x_i - x_j) = \frac{\gamma_0}{s\sqrt{2\pi}}\exp\left(\frac{-(x_i - x_j - d)^2}{s^2}\right)$$

for $x_i > x_j$. Birth function $f$ depends on species size. The second term of equation (9) is mortality rate, also depending on species size. Following Loeuille & Loreau (2005), the size dependence of the birth and mortality rates reads $f(x_i) = f_0 x_i^{-0.25}$ and $m(x_i) = m_0 x_i^{-0.25}$. Finally, the third term corresponds to competition by interference between species of similar size. Function $\beta$ is the competition by interference rate. It is a gaussian of width $s_a$ and height $b$ (competition intensity):

$$\beta(x_i - x_j) = b\exp\left(-\frac{(x_i - x_j)^2}{s_a^2}\right)$$

This model is a consumer-resource model with an abiotic compartment, but using a mass balance hypothesis, *i.e.* $\sum_{j=0} n_j = N_{tot}$, (mass conservation: Leibold, 1996), it can be, as the previous scenarios, reformulated under the Lotka-Volterra form (equation 1). Starting from the original equation (9):

$$\frac{1}{n_i}\frac{dn_i}{dt} = f(x_i)\sum_{j=0}\gamma(x_i - x_j)n_j - m(x_i) - \sum_{j=1}\beta(x_i - x_j)n_j - \sum_{j=1}\gamma(x_j - x_i)n_j$$

in the first sum, we separate the first term (j=0) from the other ones (j>0):

$$\frac{1}{n_i}\frac{dn_i}{dt} = f(x_i)(\gamma(x_i-x_0)n_0+\sum_{j=1}\gamma(x_i-x_j)n_j)-m(x_i)-\sum_{j=1}\beta(x_i-x_j)n_j-\sum_{j=1}\gamma(x_j-x_i)n_j$$

we then introduce the mass-balance constraint $n_0 = N_{tot} - \sum_{j=1} n_j$ and get, after a few rearrangements:

$$\begin{aligned}\frac{1}{n_i}\frac{dn_i}{dt} =& f(x_i)\gamma(x_i-x_0)(N_{tot}-\sum_{j=1} n_j) \\ &+ f(x_i)\sum_{j=1} n_j\gamma(x_i-x_j) - m(x_i) - \sum_{j=1}\beta(x_i-x_j)n_j - \sum_{j=1}\gamma(x_j-x_i)n_j\end{aligned}$$

we rearrange to get all the term summing over j together:

$$\begin{aligned}\frac{1}{n_i}\frac{dn_i}{dt} =& f(x_i)\gamma(x_i-x_0)N_{tot} - m(x_i) \\ &+ \sum_{j=1} n_j\left[f(x_i)(\gamma(x_i-x_j)-\gamma(x_i-x_0)) - \beta(x_i-x_j) - \gamma(x_j-x_i)\right]\end{aligned}$$

and finally we factorize by $f(x_i)\gamma(x_i-x_0)N_{tot} - m(x_i)$ and get:

$$\begin{aligned}\frac{1}{n_i}\frac{dn_i}{dt} =& (f(x_i)\gamma(x_i-x_0)N_{tot} - m(x_i)) * \\ &\left(1 - \frac{\sum_{j=1} n_j\left[f(x_i)(\gamma(x_i-x_0)-\gamma(x_i-x_j)) + \beta(x_i-x_j) + \gamma(x_j-x_i)\right]}{f(x_i)\gamma(x_i-x_0)N_{tot} - m(x_i)}\right)\end{aligned}$$

which can be recognized as a Lotka-Volterra form with

$$r(x_i) = f(x_i)\gamma(x_i-x_0)N_{tot} - m(x_i) \tag{10}$$

so that:

$$\frac{1}{n_i}\frac{dn_i}{dt} = r(x_i)\left(1 - \frac{\sum_{j=1} n_j\left[f(x_i)(\gamma(x_i-x_0)-\gamma(x_i-x_j)) + \beta(x_i-x_j) + \gamma(x_j-x_i)\right]}{r(x_i)}\right)$$

and using the hypothesis that $a_{i,i} = 1$, we have:

$$\frac{a(x_i,x_i)}{k(x_i)} = \frac{1}{k(x_i)} = \frac{f(x_i)(\gamma(x_i-x_0)-\gamma(0)) + \beta(0) + \gamma(0)}{r(x_i)}$$

so that:

$$k(x_i) = \frac{r(x_i)}{f(x_i)(\gamma(x_i - x_0) - \gamma(0)) + \beta(0) + \gamma(0)} \tag{11}$$

Equation (9) now writes:

$$\frac{1}{n_i}\frac{dn_i}{dt} = r(x_i)\left(1 - \frac{\sum_{j=1} n_j\left[f(x_i)(\gamma(x_i - x_0) - \gamma(x_i - x_j)) + \beta(x_i - x_j) + \gamma(x_j - x_i)\right]}{k(x_i)\left(f(x_i)(\gamma(x_i - x_0) - \gamma(0)) + \beta(0) + \gamma(0)\right)}\right)$$

so that function $a(x_i, x_j)$ writes:

$$a(x_i, x_j) = \frac{f(x_i)(\gamma(x_i - x_0) - \gamma(x_i - x_j)) + \beta(x_i - x_j) + \gamma(x_i - x_j)}{f(x_i)(\gamma(x_i - x_0) - \gamma(0)) + \beta(0) + \gamma(0)} \tag{12}$$

Those three functions are represented Fig. 1a in the main article. In this scenario, following Loeuille & Loreau (2005), we set $m_0 = 0.1$, $s_a = 1.5$, $\gamma_0 = 1$, $f_0 = 0.3$, $d = 2$, $x_0 = 0$ and we varied $b$ (interference intensity) and $s$ (consumption function width). Parameter values explored are given in section 2 below.

### 2 Parameter sets explored and those plotted in the main article figures

In each scenario, key parameters were varied in order to ensure that conclusions are robust to parameters changes.

- In the *Niche* scenario, the width $s_a$ of the symmetric competition function $a(x_i, x_j)$ was varied from 0.5 to 1.5. In this scenario what matters is only the ratio of $s_a$ over $s_k$, hence the two parameters need not be both varied. A value $s_a = 1$ thus means that the widths of the competition kernel and of the carrying capacity functions are identical ($s_a = s_k$).

- In the *Body-size* scenario, the width of the competition function ($s_a$) and the level of competitive asymmetry ($d$) were varied respectively from 1.2 to 2.4 and from 0.1 to 0.3.

- In the *LH-tradeoff* scenario, the preemption level ($\gamma$) and the trade-off intensity ($\alpha$) were respectively varied from 0.3 to 0.7 and from 2 to 12.

- In the *Trophic* scenario, the level of competition by interference ($b$) and the width of the consumption function ($s$) were varied respectively from 0.10 to 0.19 and from 1.0 to 1.6.

The parameter ranges presented above have been explored from the minimum to the maximum value. For clarity, in the main article, only three curves corresponding to parameter combinations yielding representative and contrasted patterns were selected and shown in figures. The parameter combinations corresponding to each curve in Figures 2-5 are provided in table S1.

| Legend | Niche | Body-size | LH-tradeoff | Trophic |
|---|---|---|---|---|
| —— | $s_a = 0.5$ | $s_a = 1.2$, $d = 0.1$ | $\alpha = 2$, $\gamma = 0.3$ | $s = 1.0$, $b = 0.10$ |
| – – – | $s_a = 0.9$ | $s_a = 1.5$, $d = 0.1$ | $\alpha = 4$, $\gamma = 0.7$ | $s = 1.0$, $b = 0.13$ |
| - - - - - | $s_a = 1.5$ | $s_a = 1.8$, $d = 0.1$ | $\alpha = 12$, $\gamma = 0.7$ | $s = 1.2$, $b = 0.10$ |

**Table S1** Parameter sets used in the main text for the four scenarios. Legend line types refers to the line types used in the figures.

### 3 Algorithm of community formation

**Co-adapted communities** We assume that each species is characterized by its mean trait value $x$ and possesses some (small) heritable variance around the latter. In these conditions, the direction and intensity of natural selection on the species trait can be determined by the selection gradient around the species trait value (Christiansen, 1991). The selection gradient is computed at the first derivative of the fitness (exponential rate of increase) of a rare variant with respect to its trait value, evaluated at the trait value of the focal species. The fitness of a rare variant with trait value $x_m$ is defined as, from equation (1):

$$F(x_m) = \lim_{n_m \to 0} \left( \frac{1}{n_m} \frac{dn_m}{dt} \right) = r(x_m) \left( 1 - \sum_j \frac{n_j a(x_m, x_j)}{k(x_m)} \right), \tag{13}$$

where all species are at their respective equilibrium abundance (Christiansen, 1991; Metz *et al.*, 1995). The selection gradient for the $i$-th species in the community is thus:

$$\nabla(x_i) = \left. \frac{dF(x_m)}{dx_m} \right|_{x_m = x_i} \tag{14}$$

If the selection gradient is positive, adaptive evolution pushes the trait value towards higher values, and if the gradient is negative, the trait moves to lower values, at a speed approximately proportional to the value of the gradient. Fitness and selection gradient are frequency- and density-dependent and change with the composition of the entire community. If the system reaches a state where the selection gradient cancels for all species, then directional evolution ceases and the community has reached a (co)evolutionary equilibrium (Christiansen, 1991; Abrams *et al.*, 1993). Note that at such an equilibrium individual species may be at an evolutionary maximum or at an evolutionary minimum. Evolutionary minima may under appropriate circumstances and inheritance modes favor the splitting of a species into two novel lineages (evolutionary branching) and thus an increase in the number of evolving lineages (Metz *et al.*, 1995; Dieckmann & Doebeli, 1999; Calcagno *et al.*, 2017). Since in this work we systematically consider all diversity levels, we can remain agnostic to second-order selection and to the specific history of transitions among diversity levels (that might involve branchings, invasions and extinctions). We just need to identify, for every diversity level, all feasible co-evolutionary equilibria, characterized by:

$$\nabla(x_i) = 0 \qquad i \in [1, \ldots, s] \tag{15}$$

For computational efficiency, for each ecological scenario and diversity level, we first identify the different coevolutionary equilibria that are attractors by iterating the adaptive dynamics of traits (defined by the selection gradient (14)) forward in time, starting from 100 random species trait values. From these we identify the one or several coevolutionary equilibria. Then, by continuation, we track these equilibria continuously through the range of parameter values, together with potential bifurcations (changes in the number or nature of equilibria). This provides an exhaustive list of coevolutionary equilibria for all parameter combinations.

**Random communities** To assemble random communities, we randomly drew each species trait value, independently, from a common probability distribution. The latter distribution can be regarded as characterizing the available regional pool of species and trait values. This species pool distribution was chosen in order to allow all feasible trait values and to be as "random" as possible: specifically, it was chosen to minimize information content, using the maximum entropy principle (Jaynes, 1957). For each scenario, we first specified the support of the distribution, *i.e.* the range of possible values for species traits. We then specified only one additional constraint, for each parameter set: the mean trait value, that had to be representative of the species typically expected for the given parameter set. Specifically, it was taken to be equal to the mean trait value $\lambda$ that would be observed in the full (saturated) community. In practice, the co-adapted communities with maximum diversity levels were used as good approximations of the saturated communities. This constraint ensured that there is no systematic (average) difference between the random species trait values and the traits that are expected in the current environment, and thus to generate communities that remain of similar nature across parameter sets. This also allows to perform comparisons between co-adapted and random communities that are not biased by the fact that species in random communities would be, intrinsically, ill-adapted. Of course, with only one constraint on the mean value, max-entropy random distributions were quite broad and all trait values had a fair chance to be picked.

In the *Body-size*, *LH-tradeoff* and *Trophic* scenarios, species trait values $x$ could take values in $]0, +\infty[$, and the entropy-maximizing distributions were thus exponential distributions of mean $\lambda$ defined as explained above. In the *Niche* scenario, trait values can spread over $]-\infty, +\infty[$ and the model is by construction symmetric around $x = 0$. Thus the random trait distribution was a double exponential, centered on zero, and with width defined by the average deviation from zero observed in the saturated community. Values for $\lambda$ are given in table S2. Remark that we also used a much more classical and simpler approach consisting in using uniform distributions for random traits, between some arbitrary defined minimum and maximum values. Results were similar, and are thus not qualitatively dependent on the precise random distributions used.

| Legend | Niche | Body-size | LH-tradeoff | Trophic |
|---|---|---|---|---|
| —— | 1.52 | 7.63 | 7.35 | 5.10 |
| – – – | 3.35 | 9.65 | 3.71 | 5.15 |
| ----- | 6.48 | 11.9 | 1.17 | 5.33 |

**Table S2** Mean trait values $\lambda$ of the saturated communities, taken to define the entropy-maximising distributions. See table S1 for the correspondence between legend line types and the parameter sets used in the main text.

In practice, for each parameter set and diversity level ($N$), random communities were formed by picking $N$ traits from the species pool defined in the previous paragraphs. The ecological equilibrium obtained from equation (1) was computed, and the community was retained (ecological filter) if all $N$ species persisted at equilibrium with a positive abundance (*i.e.* had abundance above a threshold value of $10^{-5}$). The process was repeated until 1,000 such communities were obtained. Remark that co-adapted communities (as defined in the previous section) are always a specific subset of random communities, characterized by the additional constraint (15), as illustrated in Fig. 1 of the main article.

### 4 Metrics used to quantify diversity-functioning relationships

For each BEF relationship, we computed several alternative metrics that may be used to quantify the corresponding ecosystem function. Several metrics gave identical or similar conclusions, and for clarity we retained only one or two metrics per BEF relationship that are more commonly used. A summary of all metrics can be found in Table S3 to inform the reader that no different conclusion could be drawn from those various metrics. Then, in the following, for each BEF relationships, we describe the metric(s) effectively used for the results presented in the main text.

| BEF relationship | Metrics | References |
| --- | --- | --- |
| Production | Total abundance | (1) |
| | **Productivity*** | (1) |
| Stability | **Asymptotic resilience*** | (2) (3) |
| | Stochastic invariability | (3) |
| | Initial Resilience | (3) |
| | **Ecosystem stability*** | (4) (5) |
| | Robustness | (6) |
| | Robustness heterogeneity | (6) |
| Invasion | Deterministic invasion probability | (7) (8) |
| | **Stochastic invasion probability*** | (7) (8) |
| | Proportion of the invader | (7) |
| | Mean impact of an invader on abundances | (7) |
| | **Proportion of species non extinct*** | (7) |

* *Metrics retained in the main article*

**Table S3** Metrics measured on the two types of communities for different parameter sets. Metrics in bold types are the one plotted and analyzed in the main article. Each are representative of their categories of relationships. (1) Tilman *et al.* (1996) ; (2) May (1973a) ; (3) Arnoldi *et al.* (2016) ; (4) May (1973b) ; (5) Ives *et al.* (1999) ; (6) Barabás & D'Andrea (2016) ; (7) Elton (1958) ; (8) Hector *et al.* (2001)

#### 4.1 Productivity

Community productivity is the summed productivity of each of the $N$ component species.

$$\Pi = \sum_{i=1}^{N} n_i g_i \tag{16}$$

with $g_i$ the per capita production rate of species $i$ in its community whose expression depends on the ecological interaction and community composition.

For the *Niche* and *Body-size* scenarios, there is no explicit production rate since the models are directly formulated in Lotka-Volterra form. We made the simple choice of using the intrinsic growth-rate $r_i$ as the metric per capita productivity so that :

$$\Pi_{Niche} = \sum_i n_i r_i$$

$$\Pi_{Body} = \sum_i n_i r_i$$

For the *LH-tradeoff* scenario, we consider the colonization of empty sites $(N - \sum_j n_j)$ as a contributions to productivity. This leads to the per capita productivity:

$$g_{i,LH} = n_i x_i (N - \sum_j n_j)$$

and the total productivity is the sum over all species $i$:

$$\Pi_{LH} = \sum_i n_i g_{i,LH} = \sum_i n_i x_i (N - \sum_j n_j)$$

Finally for the *Trophic* scenario, the net growth rate is straightforward, simply referring to consumption by specie $i$ of lower size species.

$$\Pi_{chain} = \sum_i n_i \left( f(x_i) \sum_{j=0} \gamma(x_i - x_j) n_j \right)$$

#### 4.2 Stability

Asymptotic resilience is taken as a measure of species stability (May, 1973a; Arnoldi *et al.*, 2016). It refers to the asymptotic return speed of the slowest species to equilibrium after an external abundance perturbation.

$$R_\infty = -\Re(\lambda_m(J)) \tag{17}$$

with $\lambda_m(J)$ the highest eigen value of the Jacobian matrix $J$ of the ecological system whose coefficient are given by (18).

$$J_{i,j} = \frac{\partial}{\partial n_j} \left( \frac{dn_i}{dt} \right)^* \tag{18}$$

To get a measure of the all community stability, we consider the community variance, namely in the variance of the sum of abundances. This measure differs from individual direction stability metrics such as the asymptotic resilience. The sum of abundances can return faster to its equilibrium value after a perturbation, even if species abundances are still fluctuating into the community. It is commonly expected (even if not general and depending on metrics) that species stability has more often a negative relationship with diversity while community stability tends to get a positive one. Mathematically,

$$var(N_T) = \sum_i var(n_i) + \sum_{i,j,j\neq i} cov(n_i, n_j)$$

with $N_T = \sum_{i=1}^{S} n_i$ the sum of each species abundance for a community with $S$ species. The variance-covariance matrix $B$ is involved into the equilibrium distribution for population fluctuation in a stochastic environment (May, 1973b) and is the solution of the Lyapunov matrix equation (19) (see also Wang *et al.* (2015) for a similar derivation):

$$D = 1/2(BA + A^T B) \tag{19}$$

with $A = N^{-1}JN$ and $N$ the diagonal matrix containing the equilibrium abundances of each species whose coefficients are $N_{i,i} = n_i^*$. $A^T$ is $A$ transpose, and $D$ the matrix containing $D_{i,j}$ coefficient which are the overall covariance between white-noise fluctuations in the stochastic differential equation of species $i$ and the one of species $j$:

$$\frac{dn_i}{dt} = r_i n_i \left(1 - \sum_j \frac{n_j a_{i,j}}{k_i}\right) + \sum_k \rho_{i,k}\gamma_k(t) n_k \tag{20}$$

with $\rho_{i,k}$ measuring the covariance between the environmental fluctuation for species $i$ and for species $j$, and $\gamma_k(t)$ is environmental stochasticity in the growth rate of population $k$ at time $t$, taken as a white noise random fluctuation with variance $\sigma^2$. $D_{i,j}$ coefficients are the diffusion coefficient involved in the Fokker-Planck equation, which is the differential equation for the probability of transition in between two states of the system. In our case, we choose to put a perturbation in a white-noise form that has no inter-species dependence: $\rho_{i,i} = 1$ ($\forall\, i$) and $\rho_{i,k} = 0$ ($\forall\, i \neq k$), and $D$ is a diagonal matrix. All individuals get the same amount of perturbation, so that each species is perturbed with an intensity that depends on its abundance. More precisely, the variance imposed on a given species is proportional to the squared of its abundance: $D_{i,i} = \sigma^2 n_i^2$. This derivation assumes sufficiently small perturbations around the equilibrium to get a local linearization of the system. To solve the continuous Lyapunov matrix equation (19), we use the *lyap()* function from Scilab. We then sum all the elements from the variance-covariance matrix $B$, and divide this sum by $\sigma^2$ (the variance of perturbation received by each individual, *e.g.* we normalize by the per individual perturbation). The coefficient of variation $CV$ is defined by:

$$CV = \frac{\sqrt{\frac{\sum_{i,j} B_{i,j}}{\sigma^2}}}{N_T} \tag{21}$$

And the community stability metrics is defined by the inverse of $CV$. Actually, in the calculation, the value of sigma has no importance because we finally divide by the same quantity. We take it equal to 1.

##### 4.3 Response to invasion

Two aspects of ecological invasion are considered: (i) the resistance to invasion and (ii) the tolerance to invasion. Resistance to invasion is defined as the probability that an alien species (randomly drawn from the regional pool and introduced at low initial abundance) does not successfully establishes in the community. The probability $P_{instal}$ that this alien species establishes is:

$$P_{instal} = \int_{x_{min}}^{x_{max}} p_x(x_e) H_{st}(x_e) dx_e \tag{22}$$

with $p_x(x_e)$ the trait distribution probability, $H_{st}(x_e) = \begin{cases} \frac{s(x_e)}{b(x_e)} & \text{if} s(x_e) > 0 \\ 0 & \text{if not} \end{cases}$, $x_e$ the trait of the foreign species which is trying to invade, $b(.)$ the growth rate and $s(.)$ the fitness function. The trait distribution probability is the same used to form the random communities (see section 3). Resistance to invasion is then defined by

$$R_{inv} = 1 - P_{instal}$$

Tolerance to invasion $T_{inv}$ (eq. 23) is defined by the proportion of species, following a successful invasion, that are not driven to extinction by the invader:

$$T_{inv} = \frac{N_{final}}{N_{init}} \tag{23}$$

with $N_{init}$ the number of species into the community right after the invasion, thus including the invasive species, and $N_{final}$ the number of species into the community once ecological equilibrium has been recovered.

##### 4.4 Metrics for trait composition

###### 4.4.1 *Trait composition representation*

We sort communities by trait values, and calculate the average trait per rank over the 1000 random communities (or few co-adapted communities when needed). For each diversity level with $N$ specie, we thus obtain $N$ random averaged trait, and $N$ co-adapted averaged trait. Values obtained for co-adapted and random traits are ploted one against the other in Fig. 3 of the main article.

###### 4.4.2 Strength of evolutionary filter

The strength of evolutionary filter is defined with trait difference in between random and co-adapted communities. We sort communities by trait values. Then, for each random community with $N$ species, we calculated the trait difference between random and co-adapted trait per rank, and we made the average over all $N$ species ranks

$$\frac{1}{N}\sum_{i=1}^{N}|x_{i,coadapt} - x_{i,random}|$$

where $i$ denotes the rank and $x_{i,coadapt}$ (resp. $x_{i,random}$) is the trait value for the $i^{th}$ co-adapted (resp.random) species. In the case of multiple co-adapted communities (*Niche* scenario), we calculated the difference to the closest co-adapted community (which is likely to be the evolutionary attractor corresponding to this random community).

###### 4.4.3 Minimum distance to the optimum trait

In the *Niche*, *Body-size*, *TF-tradeoff* scenarios, the optimum trait value $x_o$ is defined as the trait corresponding to the maximum of function $k(x)$. In the *TF trade-off* scenario, the trait value maximizing function $k(x)$ is infinite. We thus take as optimal value $x_o$ the $95^{th}$ percentile of the species pool distribution defined for the community formation. The minimum trait distance to the optimum is defined as:

$$\min_{i=1}^{N-1}(|x_i - x_o|)$$

###### 4.4.4 Average interval between species trait values

The average interval between species traits is the average two-by-two trait distance:

$$\sum_{i=1}^{N-1}\frac{|x_{i+1} - x_i|}{N-1}$$

with $x_i$ the trait of species $i$ and $N$ the number of species into the community.

### 5 Magnitude of the difference between random and co-adapted communities

In order to ascertain that co-adapted communities can be considered as different from random communities, we measured the percentile of the metric for random communities in which the mean value of the metric for co-adapted communities is (Fig. S1). In all cases, co-adapted values stand above the 8th decile or bellow the 2nd decile of the random values distribution, for at least part of the BEF relationship. This indicates that co-adapted communities are quite atypical, relative the variability within random communities.

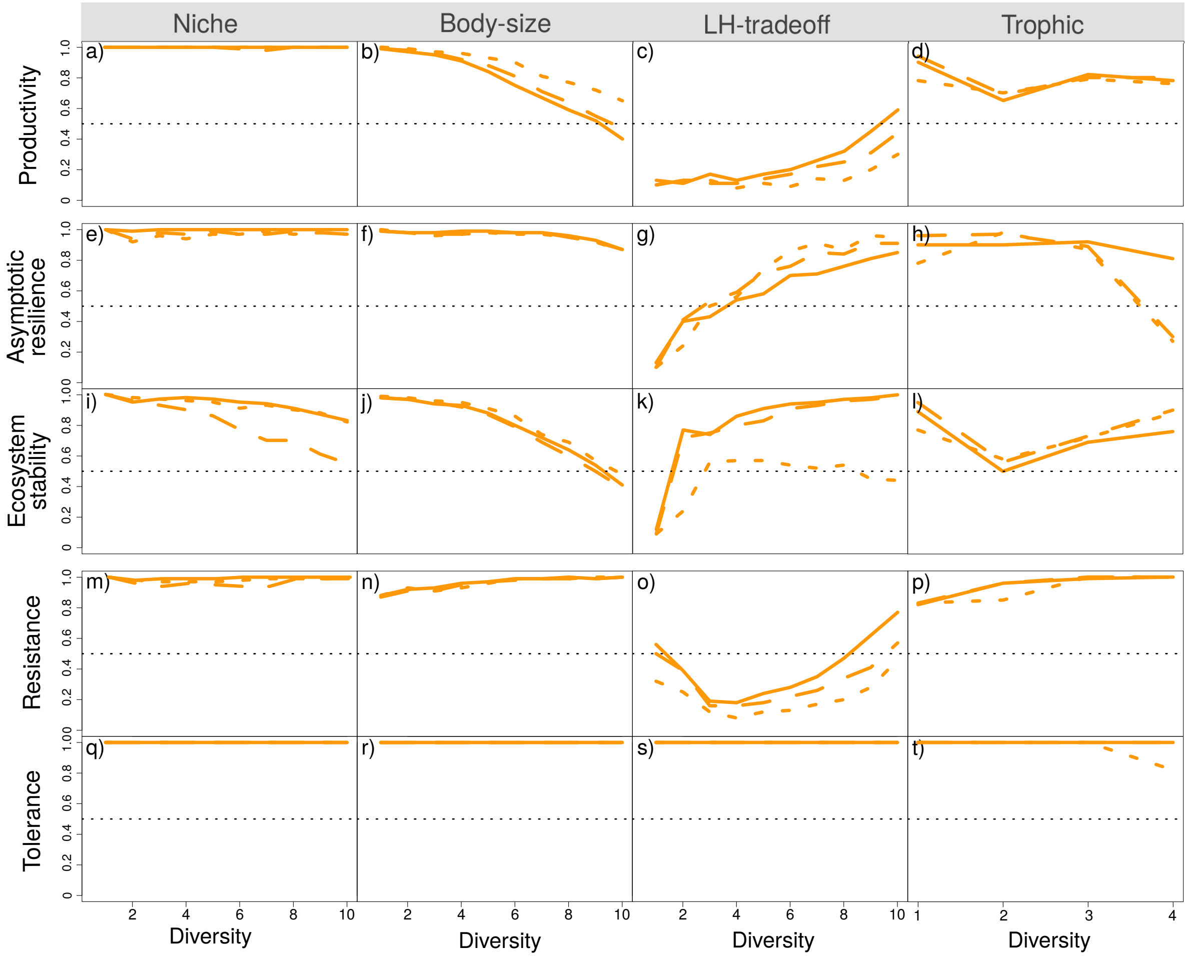

**Fig. S1** Quantile $Q_c$ of the metric for random communities in which the mean value of the metric for co-adapted communities is. It is measured for the four scenarios and 5 metrics explored (productivity a-d, asymptotic resilience e-h, ecosystem stability i-l, resistance m-p and tolerance q-t). The closest $Q_c$ to 0.5, the more similar co-adapted and random communities regarding this metric. When $Q_c$ is larger (resp. lower) than 0.5, the metric for co-adapted communities is larger (resp. lower) than for random communities. Ecological interaction parameters set are varied in each scenario (three different line types). Parameters values are given paragraph 2, together with other explored parameters sets (not shown for readability reasons).

### 6 Coefficient of variation of the average interval between trait values

In the main article, the average interval between trait values is plotted against diversity (Fig. 4e-h), but it does not give information about how evenly might be distributed traits among a particular community. To get this information, we plotted the coefficient of variation of the average interval between trait values (Fig. S2). Species traits in co-adapted communities are found more evenly distributed (lower coefficient of variation at any single diversity level) than in random communities.

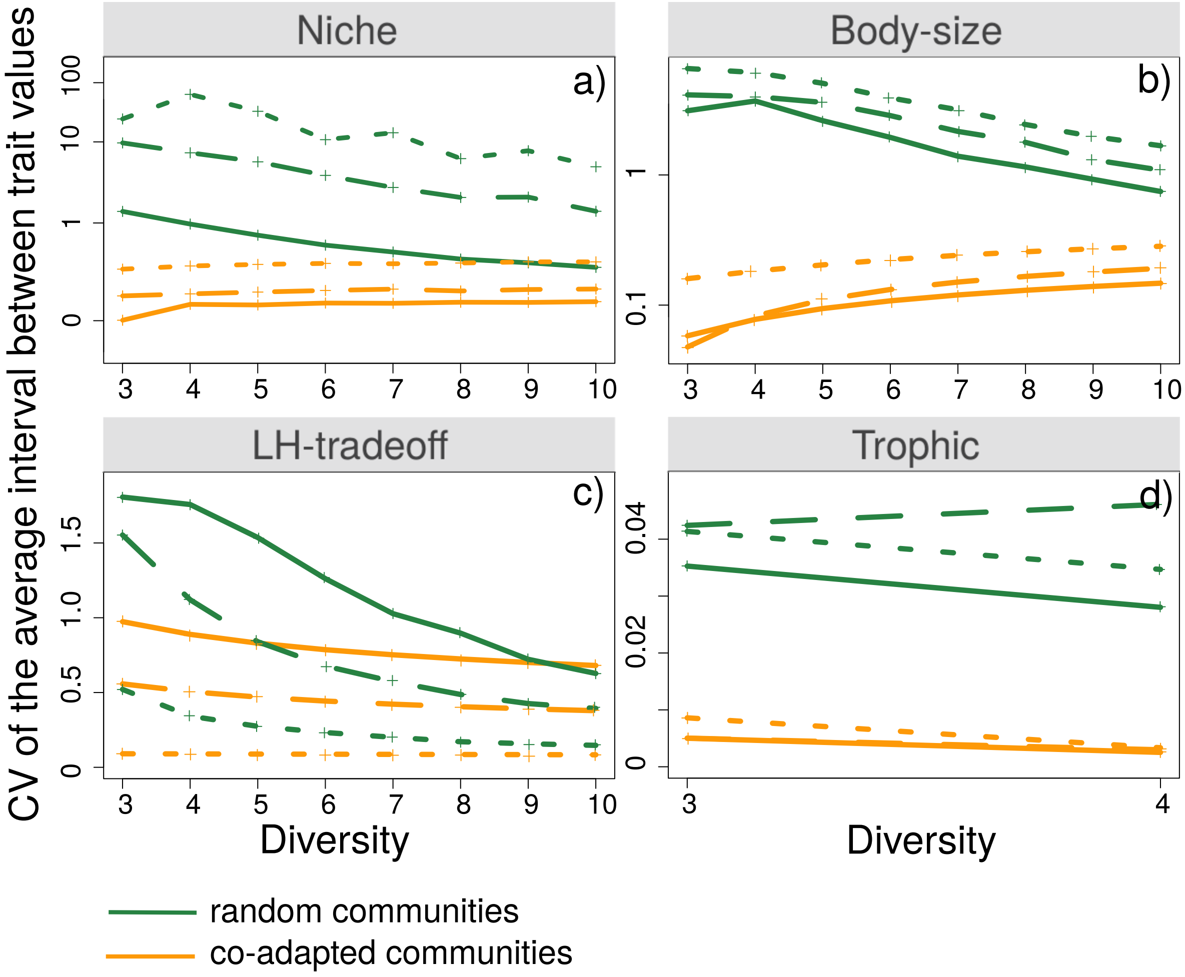

**Fig. S2** Coefficient of variation of the average interval between trait values in the four ecological scenarios. The smaller the value, the more evenly distributed are species traits. Ecological interaction parameter sets are varied in each scenario (three different line types). Parameters values are given paragraph 2, together with other explored parameters sets (not shown for readability reasons).

### 7 Results with a limit on possible trait change

The model assumes possible unbounded changes of species traits, implying either sufficient mutational supply or some kind of infinitely additive alleles. However, genetic variation is usually not infinite, and the response to selection could at some point slow down or stop. To investigate this point, we have sub-sampled the random communities, excluding the ones where at least one species would have to undergo too much trait change to reach the co-adapted state. More precisely, we have defined a maximum trait displacement $x_{max}$, which corresponds to the amount of change such that the interaction strength of evolved individuals with ancestral individuals would fall to 1%. In other words, this means that traits can only undergo changes that keep individuals in a similar 'niche' as their ancestors, in the sense that they retain non-negligible interaction strength. In practice, this also prevents a species from evolving more that the typical trait interval existing between coexisting species, in species-rich communities. Having retained only the communities with no species further from $x_{max}$, we recomputed all the metrics for the *Niche* scenario, with the new set of communities (Fig. S3).

As expected, the difference between co-adapted and random communities tends to erode. Low diversity levels are more impacted as random species were more likely to stand far from the evolutionary point (see also Fig. 3 in the main text). It follows that, for instance, the patterns for biodiversity-production relationships is less pronounced (Fig. S3f compared to Fig. S3a). Interestingly, the pattern for invasion tolerance (Fig. S3j) is not affected at all, presumably because the largest differences are observed at high diversity, where a restriction on evolutionary change has little importance. In all cases, we still observe the general trends and differences that sustain our conclusions. Even though a limit on the amount of evolutionary change allowed would weaken the reported effects, it appears that the general messages are rather robust to this.

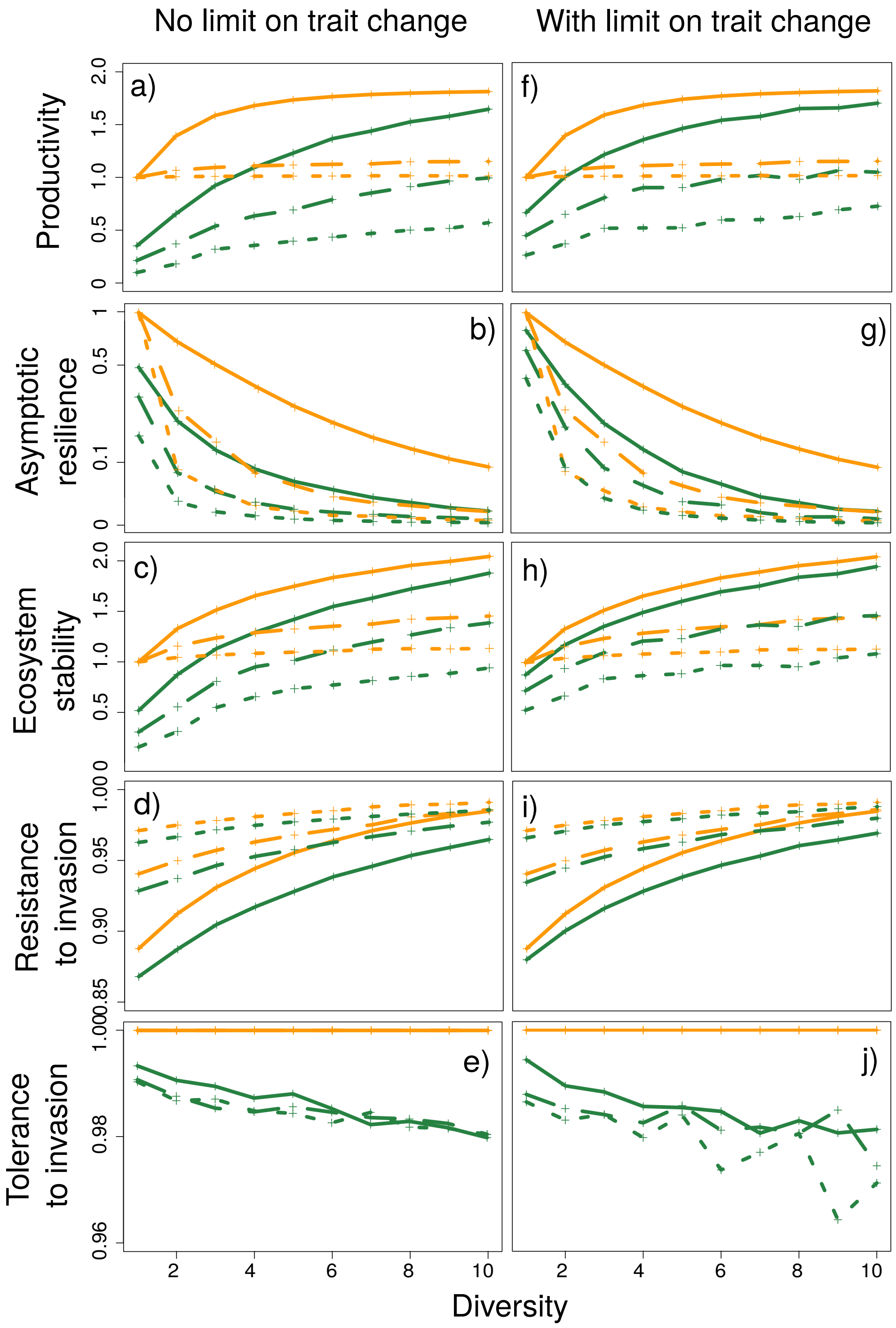

**Fig. S3** The five explored metrics for *Niche* scenario, (a-e) when all random communities are considered, and (f-j) when restricting random communities to be not too far in trait space to co-adapted ones. Parameters values (different lines type) are given paragraph 2.
